# Rank Dependency of Rescaled Pruning in Recurrent Neural Networks

**DOI:** 10.64898/2026.05.30.728930

**Authors:** Alex Q. Wang, Soon Ho Kim, Hannah Choi

**Affiliations:** Computational Science and Engineering Program, Georgia Institute of Technology, Atlanta, GA, United States; School of Mathematics, Georgia Institute of Technology, Atlanta, GA, United States

## Abstract

Throughout development and maturity, neural circuits undergo massive synaptic pruning, yielding highly sparse connectivity while preserving robust population-level computations. These population dynamics are often low-dimensional, allowing task-related computations to be formalized as trajectories within latent subspaces. How such low-dimensional dynamics are preserved amid widespread network sparsification remains unclear. Here, we investigate how different synaptic pruning rules shape low-dimensional dynamics and task performance in recurrent neural networks (RNNs). Moving beyond previous approaches focused on random sparsification of low-rank networks or networks with strictly constrained structures, we systematically evaluate how biologically motivated pruning rules interact with a network’s underlying rank. We show that post-pruning dynamics and task performance depend critically on the network’s initial rank due to distinct eigenspectral characteristics across rank regimes. Combining mathematical analysis with simulations, we demonstrate that pruning with synaptic rescaling preserves low-dimensional dynamics with minimal distortion in low-rank RNNs, but degrades in the high-rank regime. Our findings suggest that low-rank structure, combined with homeostatic synaptic rescaling, is essential for maintaining stable, low-dimensional dynamics in sparse networks.

**Author summary:** In biological neural networks, population activity is fundamentally shaped by synaptic connection strengths. During development, many of these connections are eliminated through synaptic pruning—a process thought to minimize metabolic costs and optimize computational efficiency by promoting network sparsity. However, how networks preserve their core functionalities after losing a vast numbers of connections remains an open question. Here, we combine mathematical analysis and simulations to examine how various pruning strategies affect network dynamics and computation in both low- and high-rank recurrent neural networks (RNNs). We show that uncompensated connection removal drastically alters network activity. However, implementing a homeostatic rescaling that strengthens remaining connections preserves original dynamics exclusively in low-rank networks, where connectivity is governed by a few dominant structural patterns. In contrast, pruning with homeostatic rescaling fails to maintain dynamics in high-rank networks. Our findings suggest that low-rank connectivity coupled with network homeostasis is crucial for maintaining brain function throughout significant developmental pruning.

## Introduction

Synaptic connectivity in neural circuits plays a crucial role in generating network dynamics underlying cognitive tasks [1–4], which often exhibit coordinated patterns confined to low-dimensional subspaces [5–11]. In an effort to understand the link between these low-dimensional dynamics and the network’s structural connectivity, low-rank recurrent neural networks (RNNs) have been widely studied to analytically predict trajectory geometry and stability [12–19]. However, most existing theoretical analyses of low-rank RNNs assume dense connectivity [12, 14–16], where every neuron connects to all others. This assumption contradicts the biological observation that neural networks observed in the brain are highly sparse and typically full-rank [20–25].

Such sparse, high-rank networks— observed across systems ranging from cortical networks [22, 26, 27] to fly connectome [28]— can emerge from large-scale synaptic pruning. Neural systems undergo extensive synaptic pruning during early development, followed by activity-based refinement across the lifespan, which eliminates a substantial fraction of initially formed connections [29–33]. While pruning reduces metabolic and physical wiring costs [34–36], it may also constrain computational capacity by reducing the number of adjustable parameters within the network [37–39]. Yet, despite this substantial loss of degrees of freedom from synaptic pruning, neural circuits continue to exhibit robust, low-dimensional population dynamics during cognitive tasks.

Sparse RNNs that preserve task-relevant, low-dimensional dynamics can be obtained through supervised learning or *post hoc* fine-tuning, where task-driven optimization identifies effective sparse subnetworks [40–44]. In contrast, for more biologically plausible unsupervised pruning where connections are removed without task-specific training, there is currently no general method for preserving recurrent dynamics. Recently, a few efforts have begun to describe how unsupervised pruning affects RNN dynamics [20, 45]. However, predictions made by the previous studies often require restrictive constraints on network structure or dynamics. For example, Moore and Chaudhuri [45] showed that noise-correlation-based pruning can preserve network dynamics in diagonally-dominant RNNs with linear dynamics; however, this approach does not extend to nonlinear regimes. Herbert and Ostojic [20], on the other hand, demonstrated that simple, random edge removal in low-rank RNNs leads to sparsity-dependent distortions of low-dimensional trajectories unless the outlier eigenvalue location is explicitly modulated, highlighting the challenge of unsupervised pruning in nonlinear recurrent networks. Their analysis was limited to random sparsification of low-rank RNNs, leaving the distinct effects of different pruning rules on both low- and high-rank RNNs unexplored.

Here, we examine how different synaptic pruning methods alter RNN dynamics across network ranks. Specifically, we focus on the role of synaptic rescaling in preserving the post-pruning dynamics. We propose *rescaled pruning*, a method in which connections are removed according to either constant or weight-dependent probabilities, while the remaining synaptic weights are strengthened to maintain the average connectivity strength. This implementation of synaptic rescaling during pruning mirrors homeostatic synaptic scaling, a biological process where neurons globally scale synaptic strengths to stabilize firing rates despite connectivity perturbations [46–60]. We systematically evaluate the impact of rescaled pruning in both low- and high-rank RNNs. In the low-rank case, the clear separation between outlier and bulk eigenvalues allows rescaled pruning to preserve low-dimensional trajectories with minimal distortion, thereby maintaining the network’s ability to perform stable computations. In contrast, the same procedure fails in the high-rank regime. We show that this failure is driven by an expansion of the bulk eigenspectrum following rescaling, combined with a pruning-induced rotation of the singular vectors that are not corrected by rescaling. Together, these factors disrupt the RNN dynamics in the high-rank regime. Our results demonstrate that pruning preserves low-dimensional trajectories in RNNs only when the original network is low-rank and accompanied by synaptic rescaling, revealing a critical rank dependency in homeostatic, pruned network dynamics.

## Results

### Pruning Rules on RNNs with Different Ranks

#### RNN model

We primarily consider non-linear RNNs of the form

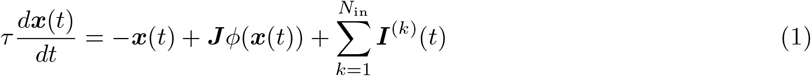

where ***x***(*t*) ∈ ℝ^*N*×1^ denotes the firing rates of *N* neurons at time *t*, with its component *x*_*i*_(*t*) representing the firing rate of the *i*-th neuron. The connectivity matrix ***J*** ∈ ℝ^*N*×*N*^ specifies recurrent connections within the network, with *J*_*ij*_ indicating the connection strength from neuron *j* to neuron *i*. The RNN receives *N*_in_ input vectors ***I***^(*k*)^(*t*) ∈ ℝ^*N*×1^ over time. The term −***x***(*t*) captures the intrinsic leaky dynamics of neuronal activity, characterized by the time constant *τ* . The activation function *ϕ*(·) introduces nonlinearity. Here *ϕ*(·) = tanh(·).

The RNN has *N*_out_ output channels, with the *l*-th readout *z*_l_ at time *t* defined as

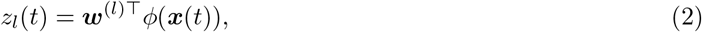

where ***w***^(l)^ ∈ ℝ^*N*×1^ denotes the output weight for the *l*-th output channel.

The rank of the connectivity matrix prior to pruning is crucial for our analysis. The connectivity matrix ***J*** can be expressed as a sum of unit rank forms,

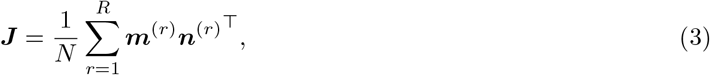

where *R* denotes the rank of ***J*** and *R* ≪ *N* . Here, ***m***^(*r*)^, ***n***^(*r*)^ ∈ ℝ^*N*×1^ denote column vectors.

#### Rank-Constrained Training

We train RNNs using rank-constrained backpropagation. Specifically, the initial recurrent connectivity matrix ***J*** is sampled from a Gaussian distribution *J*_*ij*_ ∼ 𝒩 (0, 1*/N* ). During training, after each epoch, we perform singular value decomposition (SVD) on the updated weight matrix ***J*** ^⋆^ and truncate it to the first *R* rank components. Such a procedure guarantees that the trained connectivity ***J*** is of rank of *R*. Our training process is detailed in Algorithm 1.

##### Algorithm 1

Rank-Constrained Backpropagation

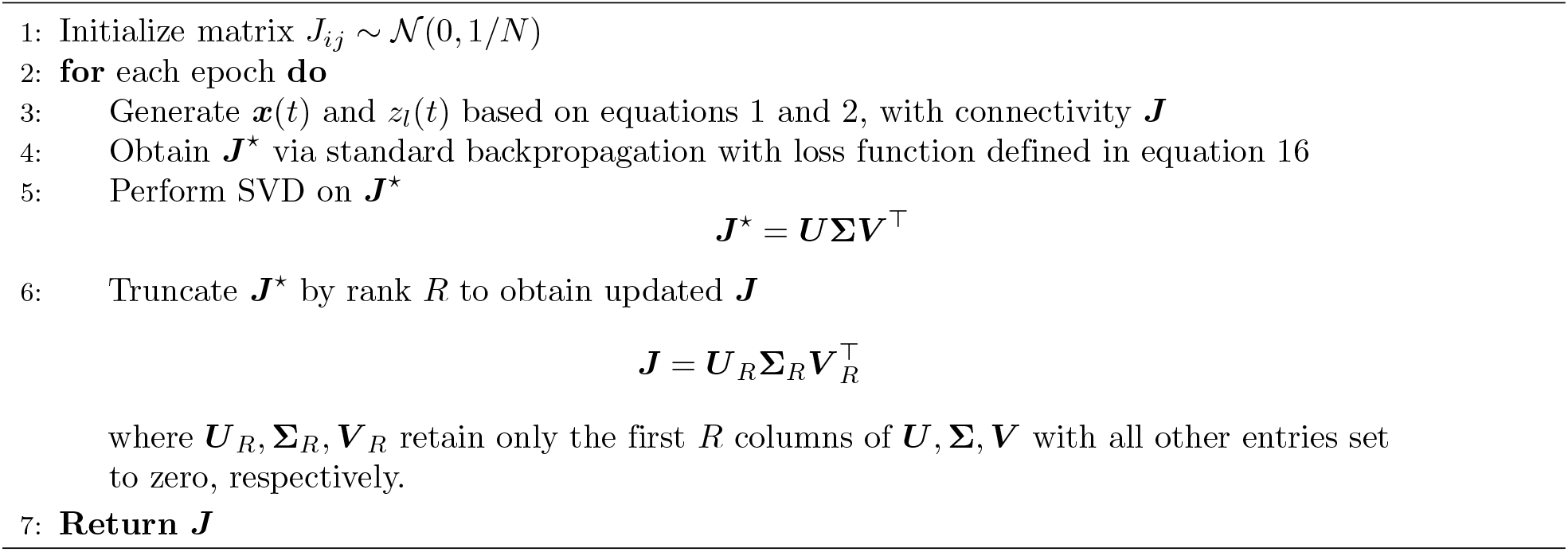

#### Pruning Rules

The goal is to use pruning to generate a sparse connectivity matrix 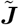 from the original connectivity matrix ***J*** while preserving the essential dynamics of the original RNN. Here, we consider unsupervised pruning rules, where each connection *J*_*ij*_ is retained independently with probability *p*_*ij*_. This can be written as applying a mask ***X*** to the connectivity matrix ***J***, resulting in the sparsified matrix:

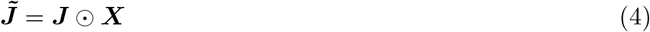

We compare two different methods of assigning weights to the remaining connections. In both methods, edges are removed or retained according to the probability *p*_*ij*_, but the scaling of the retained connection strengths differs. In the first case, which we call rescaled pruning, the remaining synaptic weights are strengthened to preserve the overall connectivity strength after pruning. This rule is analogous to biological homeostatic synaptic scaling, where neurons globally scale synaptic strengths to stabilize firing rates [46–60]. Our implementation aligns with a previous modeling approach for homeostatic synaptic scaling in which the total synaptic weight is kept fixed [61]. In our study, rescaled pruning is defined as

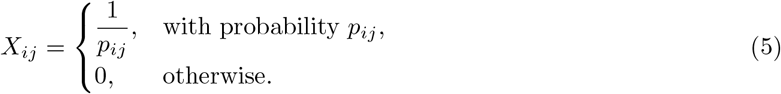

For comparison, we consider a simple pruning counterpart, where the remaining connectivity is not rescaled. Simple pruning is defined as

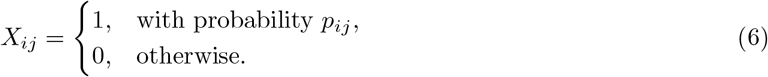

While rescaled and simple pruning dictate how retained connections are scaled, we also evaluate distinct probability criteria for selecting edges to retain. In this study, we explore two choices of *p*_*ij*_:

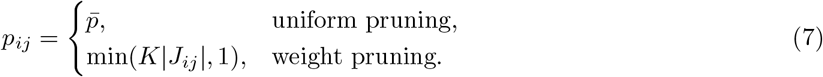

In uniform pruning, the probability *p*_*ij*_ of preserving the connection from neuron *j* to *i* is controlled by the scalar-valued parameter 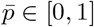 which represents the uniform probability of preserving the connection. In weight pruning, *p*_*ij*_ is controlled by the scalar-valued parameter *K* ≥ 0 and the connectivity weight *J*_*ij*_, where *K* controls the overall connection density. For both pruning models, we use *d* to represent the density of the connectivity matrix after pruning, i.e.,

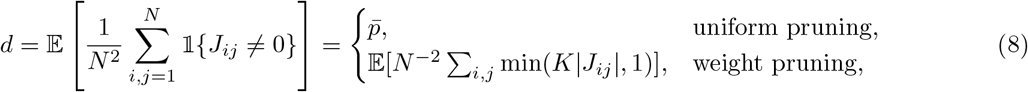

where 𝟙 is the indicator function.

### Rescaled Pruning Preserves Hidden Dynamics in Rank-One RNNs

We demonstrate that, when the connectivity matrix ***J*** is of rank-one, pruning with rescaling effectively preserves the eigenvalue spectrum of the connectivity, thereby maintaining the original low-dimensional RNN dynamics.

Specifically, for strictly rank-one connectivity, the dynamics of the network are constrained to a two-dimensional latent space spanned by vectors ***m*** and ***I*** [12]. Although pruning formally transforms 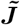 into a full-rank matrix, its effective dynamics remain dominated by a single leading real eigenvalue as a spectral outlier, which is accompanied by a bulk of eigenvalues around origin introduced by pruning. Rescaling preserves the leading eigenvalue, thus ensuring that the hidden dynamics of the pruned RNN are largely maintained despite the apparent increase in rank.

#### Eigenvalue Spectrum

We first present analytical results on the eigenvalue spectrum. Consider a rank-one connectivity matrix defined as 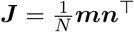 with ***m, n*** ∈ ℝ^*N*×1^, where each pair of entries *m*_*i*_ and *n*_*i*_ are drawn from a joint Gaussian distribution characterized by:

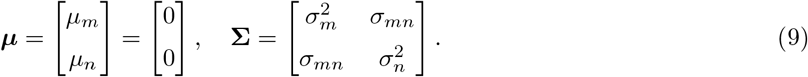

By construction, the connectivity matrix *J* only has one non-zero eigenvalue given by 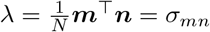 in the large *N* limit.

After pruning, the eigenvalue spectrum of the sparse connectivity matrix 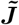 consists of two distinct components: (1) a bulk of complex eigenvalues centered around the origin, arising from the randomness introduced by pruning, which follow Girko’s circular law [62, 63], and (2) a leading real eigenvalue positioned outside this bulk, reflecting the original rank-one structure (Fig 1A and B). Table 1 summarizes the analytical results for the leading eigenvalue and the radius of the bulk spectrum after uniform or weight pruning with or without rescaling, for rank-one RNNs. The full derivation is shown in Methods.

**Table 1.** Spectral quantities of a rank-one connectivity matrix under different pruning schemes. We compare the leading eigenvalue and bulk radius *r*_*b*_ across uniform and weight pruning, both with and without rescaling. For weight pruning, the scaling factor is given by 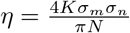.

| Quantity | Uniform Pruning |  | Weight Pruning |  |
| --- | --- | --- | --- | --- |
|  | Rescaled | Simple | Rescaled | Simple |
| Leading Eigenvalue | $\lambda$ | $\bar{p}\lambda$ | $\lambda$ | $\eta\lambda$ |
| Bulk Radius $r_b$ | $\sigma_m\sigma_n\sqrt{\frac{1-\bar{p}}{\bar{p}N}}$ | $\sigma_m\sigma_n\sqrt{\frac{\bar{p}(1-\bar{p})}{N}}$ | $\sqrt{\frac{2\sigma_m\sigma_n}{K\pi} - \frac{\sigma_m^2\sigma_n^2}{N}}$ | $\sqrt{\frac{\eta^2\sigma_m^2\sigma_n^2}{N} + \frac{8K(1-2\eta)\sigma_m^3\sigma_n^3}{\pi N^2}}$ |

**Fig 1.**
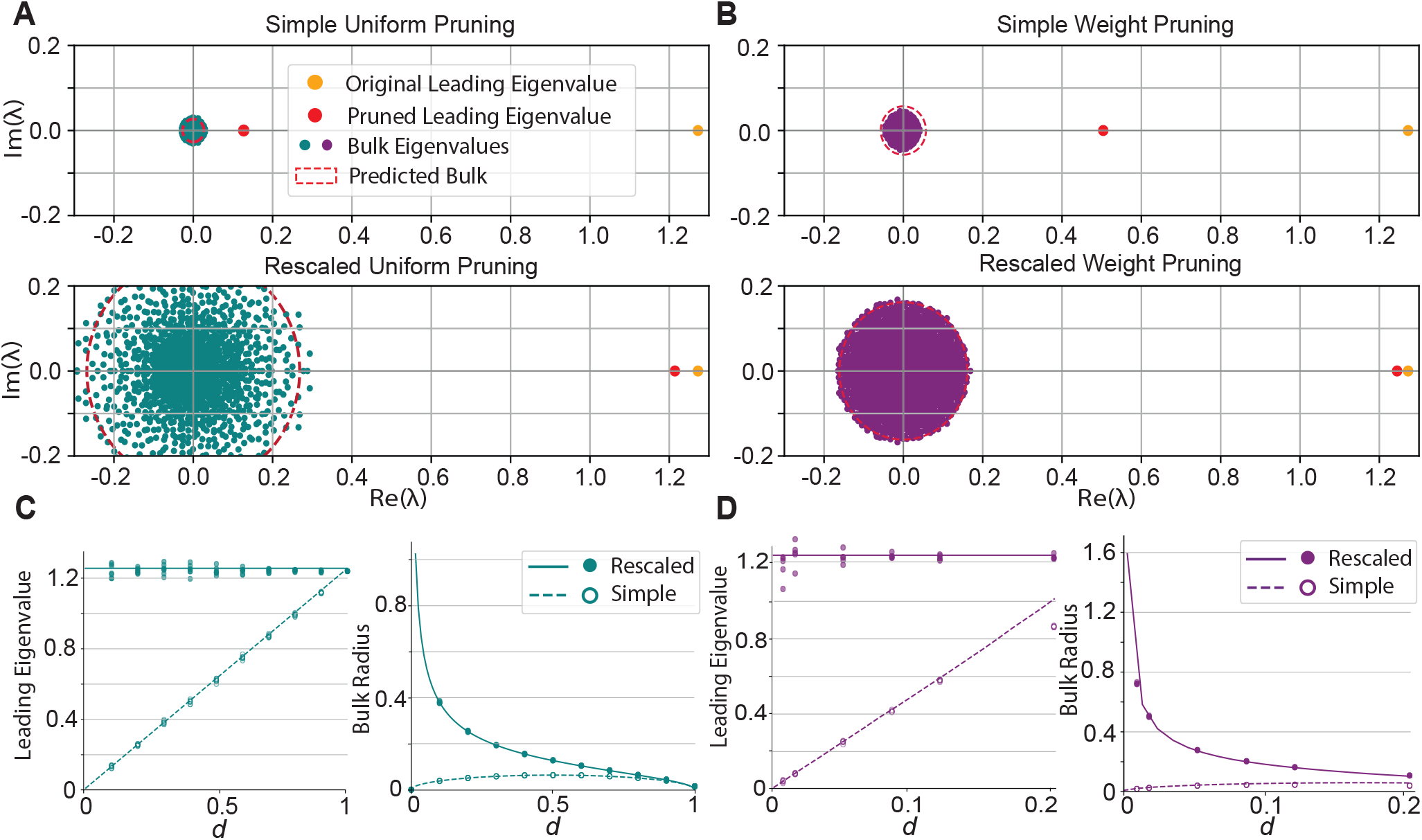
Properties of the eigenvalue spectrum of rank-one RNNs. Turquoise denotes uniform pruning, while purple denotes weight pruning. **A, B**: Eigenvalue spectrum of a random rank-one RNN pruned to density *d* = 0.4. The orange dot indicates the original leading eigenvalue of the RNN before pruning. The red dot marks the leading eigenvalue after pruning, and the purple and turquoise dots represent bulk eigenvalues introduced during pruning. The red dashed line shows the theoretical bulk radius. **C, D**: Dependence of the leading eigenvalue and the bulk radius on density *d*, respectively. Lines indicate the theoretical estimates (Table 1), and dots indicate experimental results over 5 pruning trials. The experimental bulk radius is measured as the average of the 50 largest eigenvalues in an RNN of size *N* = 1000.

In the large *N* limit, rescaled pruning consistently preserves the original eigenvalue. In contrast, despite producing a relatively smaller bulk, simple pruning reduces the magnitude of the leading eigenvalue (Fig 1A and B). Notably, the magnitude of this distortion is inversely proportional to the connectivity density *d* (Fig 1C and D).

#### Trajectory Preservation

We further verify the effectiveness of rescaling by comparing neural trajectories in latent space. In the random rank-one RNN scenario, vectors ***m*** and ***n*** are drawn from the previously defined joint Gaussian distribution, and we set the external input vector ***I*** = ***n*** to ensure the two-dimensional trajectory [12] (Fig 2A). The latent dynamics are visualized by projecting neural activities onto ***m*** and ***I***.

**Fig 2.**
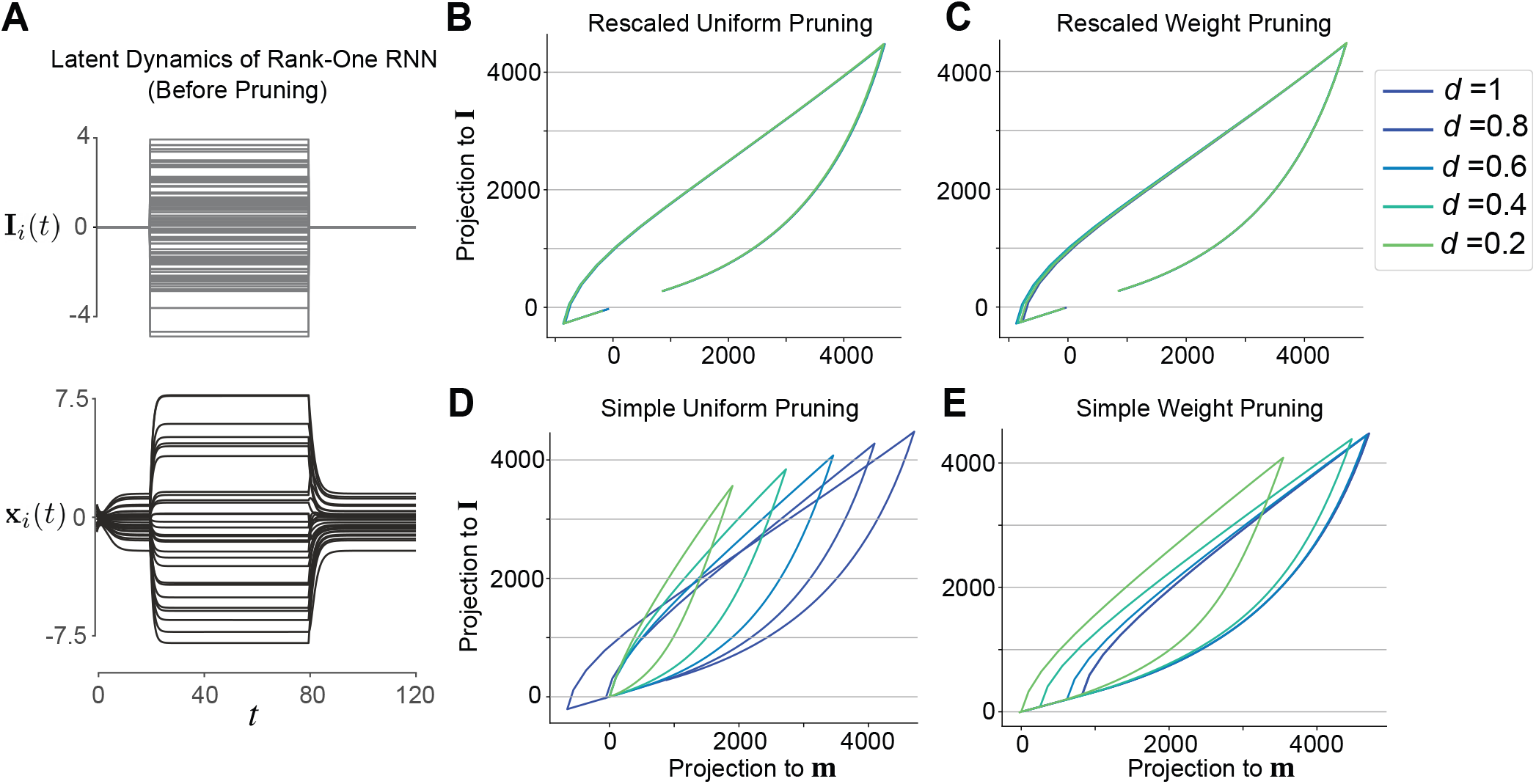
Pruning on random rank-one RNNs. **A**: Input ***I*** and hidden state ***x*** over time in a rank-one RNN before pruning, shown for 25 representative neurons out of *N* = 800. **B–E**: Hidden trajectories projected on vectors ***m*** and ***I*** of the pruned random rank-one RNN of different density levels *d*. Note that rescaled pruning always preserves hidden dynamics.

Across both random pruning and weight pruning conditions, we observe that rescaling effectively preserves the original low-dimensional dynamics, maintaining stable trajectories even as the connectivity density *d* decreases (Fig 2B and C). In contrast, when rescaling is omitted, simple pruning consistently induced distortions of the latent dynamics (Fig 2D and E). Notably, the magnitude of these distortions diminish gradually with higher connectivity densities *d*, highlighting that simple pruning is particularly unreliable in sparse RNNs. These findings reinforce the view that rescaling acts as a stabilizing mechanism, compensating for connectivity loss introduced by pruning, thus enables the network to preserve its low-dimensional dynamics.

We next extend our results to task-trained RNNs. Here, the RNN is trained using rank-constrained backpropagation (Algorithm 1) to ensure the connectivity ***J*** is of rank one. We primarily study a flip-flop task in the form of a common behavioral paradigm [64]. The network receives input pulses arriving at random intervals with *N*_in_ input channels and output channels. Upon receiving a pulse at a specific input channel, the corresponding output channel transitions to a steady-state value determined by the input signal, while all other output channels settle to zero. The network must maintain this output configuration until the next input pulse occurs. In this section, we set *N*_in_ = 1 (Fig 3A).

**Fig 3.**
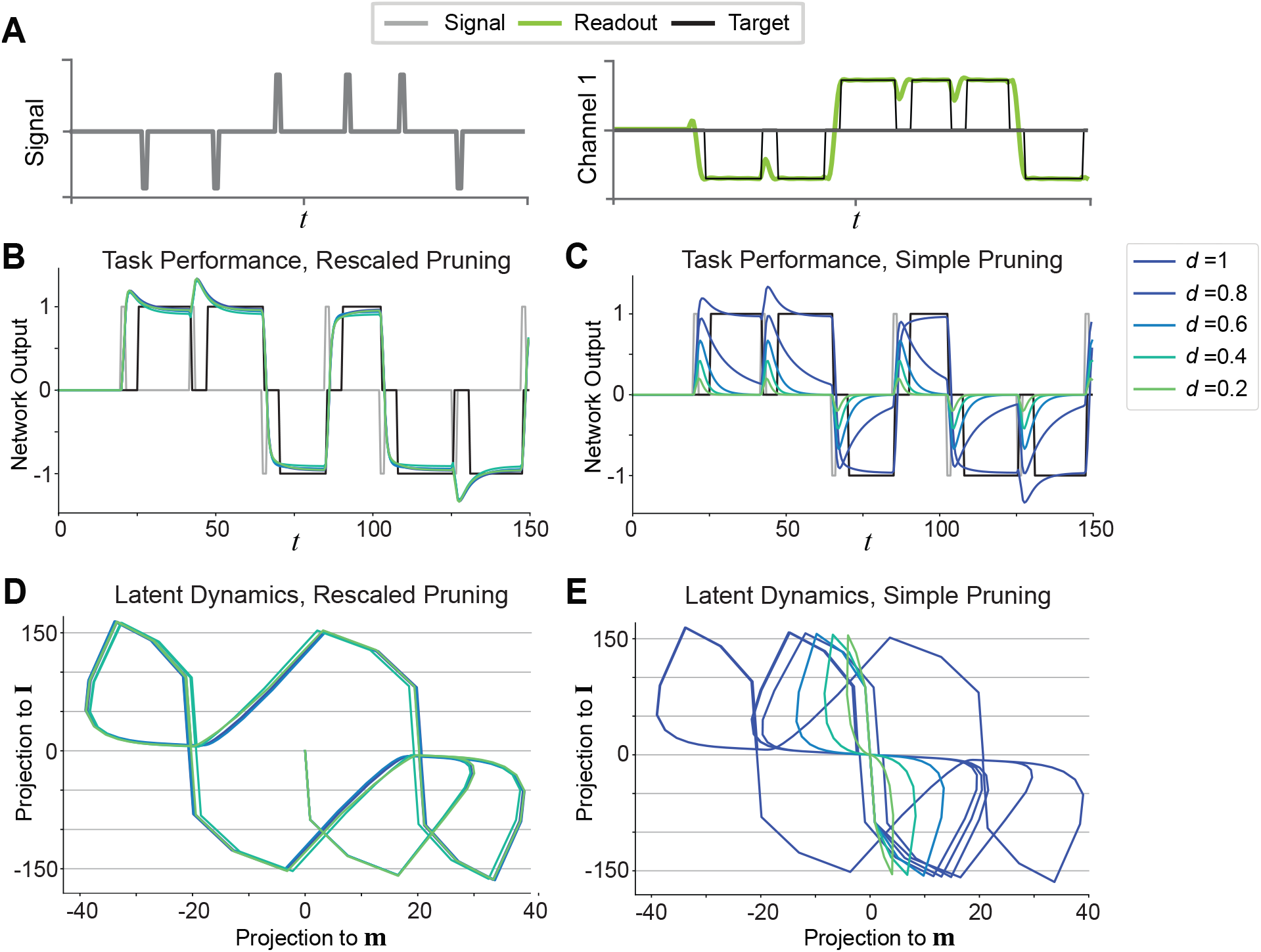
Pruning in task-trained rank-one RNNs. **A**: Schematic of a single channel flip-flop task. The left panel shows the signal delivered to the RNN; the right panel displays the target and actual readouts after training. **B, C**: Task performance of the single-channel flip-flop task in a rank-one RNN across different density levels *d* with rescaled (**B**) and simple (**C**) uniform pruning. The RNN consists of *N* = 500 neurons. Corresponding results for weight pruning are not shown but are similar. **D, E**: Hidden trajectories of the pruned rank-one RNNs under rescaled (**D**) and simple (**E**) pruning.

We show that rescaled pruning preserves task-relevant neural trajectories across a broad range of connectivity densities, thereby maintaining high task accuracy (Fig 3B and D). This robustness indicates that strengthening the remaining synaptic weights effectively compensates for the reduced number of connections, allowing the recurrent dynamics to retain the low-dimensional structure necessary for reliable computation. In contrast, simple pruning, in which connections are removed without rescaling, produced substantial distortions of the latent trajectories and led to degraded behavioral performance (Fig 3C and E). The divergence between rescaled and simple pruning is consistent with the observations in random rank-one RNNs (Fig 2), where rescaled pruning consistently preserves the dynamics, while simple pruning introduces distortions.

Taken together, these findings suggest that in rank-one RNNs, while sparsity alone can be detrimental to network function, rescaling provides a homeostatic mechanism that stabilizes dynamics and preserves hidden trajectories and computational capacity.

### Rescaled Pruning Performance Declines with Added Ranks

In previous sections, we focus on pruning in rank-one RNNs, where neural dynamics is constrained to a two-dimensional subspace. However, rank one is a restrictive case and cannot support the richer dynamics required for more complex cognitive tasks [15]. To model more realistic neural computations, we extend our analysis to RNNs with higher-rank connectivity.

#### Quantitative Evaluation

We first conduct a quantitative analysis on randomly connected high-rank RNNs following Eq 3, in which each pair (***m***^(*r*)^, ***n***^(*r*)^) ∼ 𝒩(***µ*, Σ**) constructs a unit-rank component. We construct a series of random rank-*R* RNNs according to Eqs 3 with 9 and *σ*_*m*_ = *σ*_*n*_ = *σ*_*mn*_ = 1. Because we allow *R* to vary up to 500, where the *R* ≪ *N* assumption breaks down, we scale each connectivity matrix by dividing it by the absolute value of its largest eigenvalue, so that the absolute value of the largest eigenvalue is 1 independent of the rank R (however, see Methods for a modification of Eq 3 that preserves the largest eigenvalue).

The dynamical variables ***x*** are initialized by drawing from a normal distribution. A random step input is applied to the network from *t* = 30-90 (an example of the input and network response is shown in Fig 4A). We measure the relative error between the pruned and original trajectories over time across different connectivity ranks, defined as

**Fig 4.**
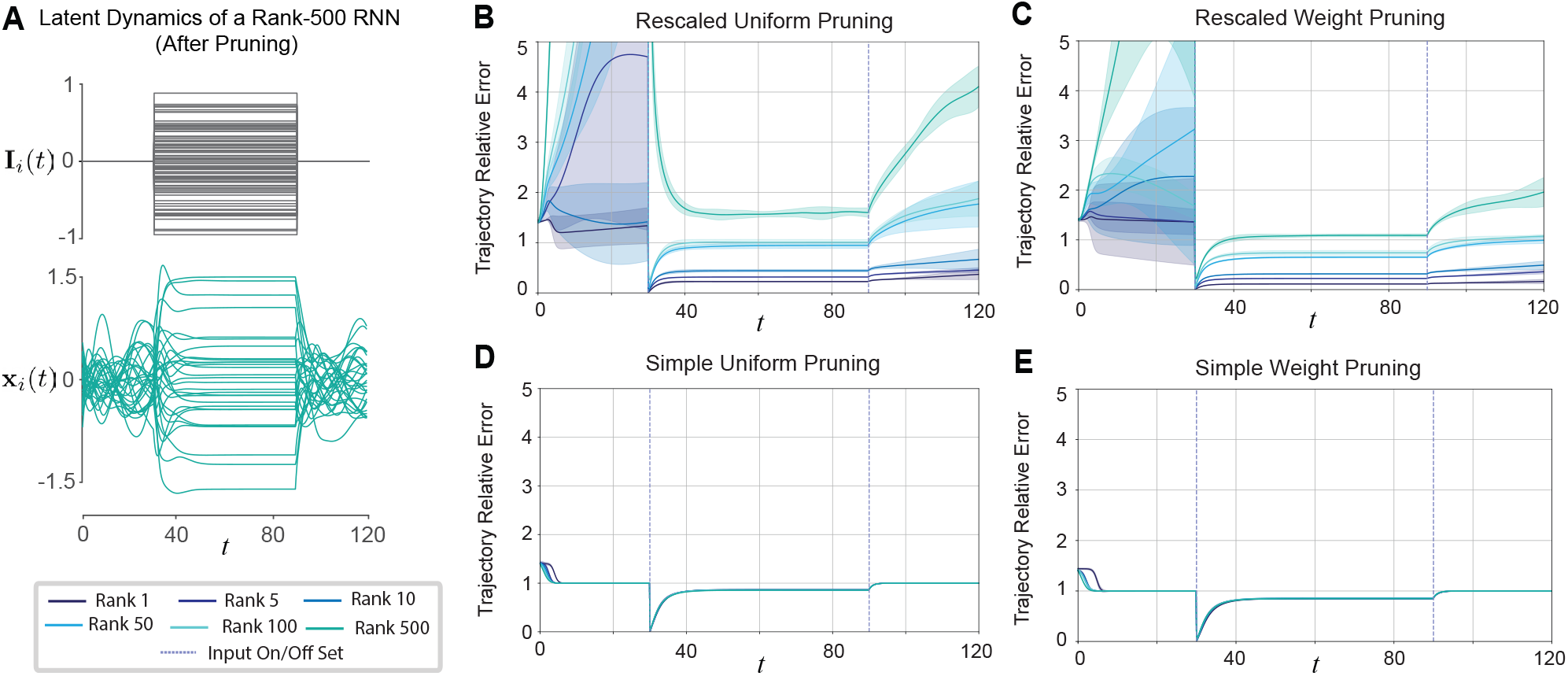
Trajectory relative error introduced by pruning in RNNs across different ranks. **A**: Input ***I*** and hidden state ***x*** for post-pruning RNN over time, shown for 30 representative neurons out of *N* = 800. **B-E**: Trajectory relative errors over time for RNNs with different ranks. The RNN is pruned to the density *d* = 0.05. The dashed lines indicate the input-driven regime, and the shaded area denotes the 1*σ* standard deviation of trajectory relative error over 5 pruning trials across time. Note that higher ranks often lead to larger trajectory errors.

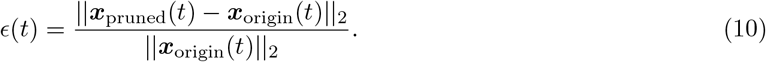

This allows us to examine how pruning affects neural trajectories under both input-driven and input-free conditions.

Our results show that increasing the rank of the connectivity matrix often leads to larger trajectory errors, reflecting a greater distortion in the underlying dynamics. This trend is observed in both random uniform pruning and weight pruning conditions (Fig 4B-E). Interestingly, in the high-rank regime, simple pruning (Fig 4D and E) yields smaller errors than rescaled pruning (Fig 4B and C), indicating that the compensatory effect of rescaling diminishes as rank increases. These observations suggest that while rescaling is beneficial in preserving low-dimensional dynamics in lower rank networks, its effectiveness is rather limited in high-rank networks, where the low-dimensional dynamics is supported by a richer set of modes.

To evaluate task performance after pruning across ranks, we measure the mean squared error (MSE) of the readout before and after pruning in rank-constrained RNNs trained on single- and double-channel flip-flop tasks (Fig 5A). Pruning primarily alters the singular value spectrum by introducing smaller noise singular values at higher ranks, while the dominant rank-*R* structure remains largely preserved (Fig 5B). To account for the inherent flexibility of recurrent networks, the output weights ***w***^(*l*)^ are retrained after pruning, thus ensuring that performance changes primarily reflected alterations in the recurrent dynamics rather than limitations of the readout weights.

**Fig 5.**
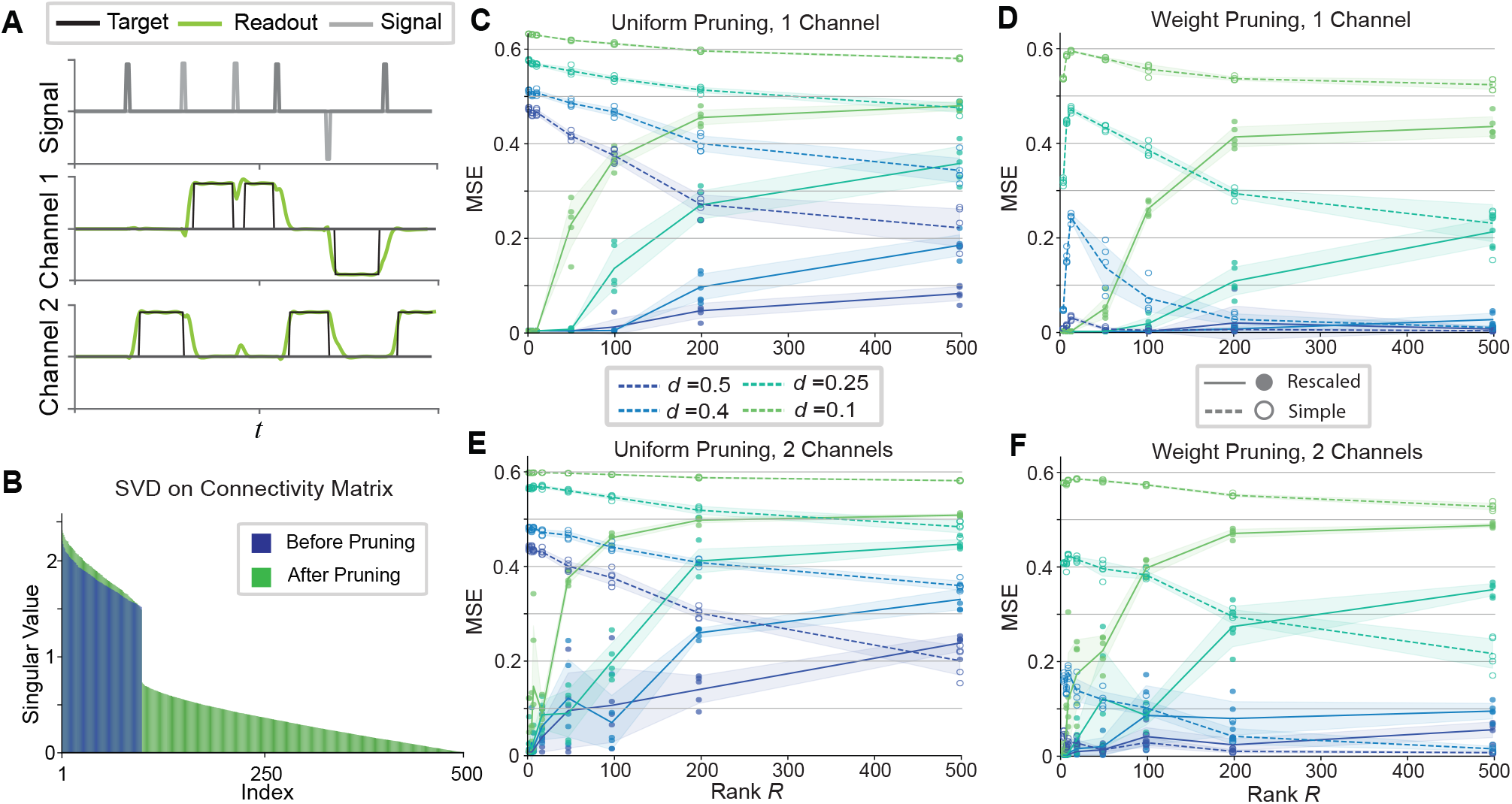
Error introduced by pruning in task-trained high-rank RNNs. **A**: Schematic of a double-channel flip-flop task. **B**: Singular value decomposition of the connectivity matrix from rank-constrained backpropagation training. The target rank is 100 with a total of *N* = 500 neurons. Blue and green bars show the singular value distribution before and after pruning, respectively. **C-F**: Mean squared error (MSE) for the single- and double-channel flip-flop task, from RNNs of different ranks with readout retraining. Open circles indicate the MSE for each of 7 trials of simple pruning, while filled circles indicate the MSE of rescaled pruning. Solid and dashed lines show the trial averages of rescaled and simple pruning, respectively. Shaded regions represent the 1*σ* deviation. Note that MSE increases with rank under rescaled pruning, while the trend is reversed under simple pruning. Increased sparsity induces higher MSE across all pruning methods.

Consistent with the observations in dynamics, the MSE under rescaled pruning increases monotonically with the rank of pre-pruning connectivity ***J*** . Notably, these results suggest the existence of a threshold in rank above which pruning introduces significant errors, reflecting a transition between regimes in which low-dimensional dynamics can be preserved and those in which pruning irreversibly disrupts dynamics relevant for computation (Fig 5C-F, solid lines).

The effects of retraining further reveals a divergence between pruning schemes. In the case of simple pruning, retraining the readout vectors results in lower MSE in high-rank RNNs compared to low-rank ones, indicating that output re-optimization can partially compensate for distortions in the recurrent dynamics (Fig 5C-F, dashed lines). By contrast, in the rescaled pruning condition, MSE continues to increase with rank despite retraining (Fig 5C-F, solid lines). This pattern suggests that while rescaling preserves the network dynamics of low rank RNNs, it may constrain flexibility at higher ranks. One possible explanation is that rescaling amplifies contributions from surviving connections, thereby limiting the capacity of retraining to realign outputs with the task-relevant subspace. In contrast, simple pruning, while generating more distorted dynamics, preserves a more balanced distribution of contributions across connections, enabling greater adaptability when readout vectors are retrained.

The decreased performance with higher rank may therefore be explained by shifts in the eigenvalue spectrum with pruning that our rescaling rule cannot preserve. Another factor that should be considered is the effect of pruning on the left and right singular vectors, which determine network response to input and its output [12]. Even if the eigenvalue spectrum is perfectly preserved, the dynamics will lie in a different manifold if the singular vectors are distorted. In order to understand the rank-dependence of RNN performance under both simple and rescaled pruning, we examine both factors.

#### Eigenvalue Spectrum

In low-rank RNNs, the dynamics are dominated by one single leading eigenvalue, which can still be effectively preserved through rescaling. However, in high-rank cases, the eigenvalue spectrum becomes too disordered to meaningfully identify a dominant mode. Instead of a single leading eigenvalue driving the dynamics, distributed eigenvalues collectively contribute to low-dimensional dynamics, making it difficult to retain all of them after pruning.

The failure of rescaled pruning in high-rank regimes may be rooted in chaotic dynamics. Since ***J*** is initialized as a Gaussian random matrix (*J*_*ij*_ ∼ 𝒩 (0, *σ*^2^) with *σ* = 1) before being task-trained, it can be still treated as a random matrix following the same distribution after rank-constrained training where the network preserves a sufficiently high rank, which, by Girko’s circular law [62, 63], makes the spectrum a bulk. Table 2 gives us the theoretical bulk radius of the eigenvalue spectrum of a full-rank RNN that is pruned either with or without rescaling, following both uniform and weight pruning. For the full derivations, see Methods.

**Table 2.**
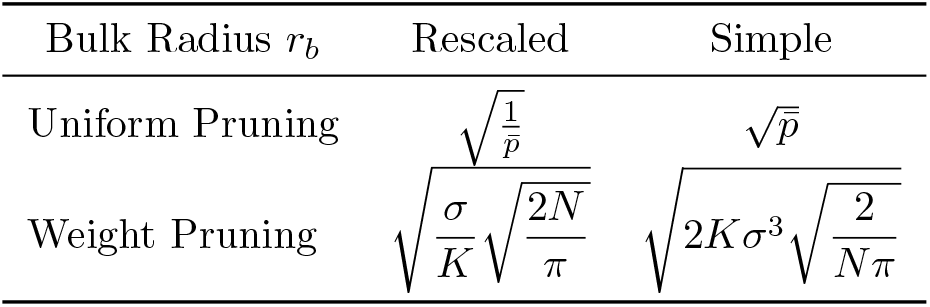
Bulk radius *r*_*b*_ of the eigenvalue spectrum under rescaled and simple pruning on a full-rank RNN with its connectivity matrix entries drawn from a Gaussian distribution, *J*_*ij*_ ∼ 𝒩 (0, *σ*^2^), for both uniform and weight pruning schemes.

Notably, rescaled pruning tends to inflate the spectral bulk of the connectivity matrix (Fig 6A and B). As the bulk radius *r*_*b*_ exceeds 1, the system transitions into a chaotic regime, in which trajectories become unstable and the network loses its capacity to sustain task-relevant dynamics (Fig 4A, input-free regime) [65, 66]. In contrast, simple pruning reduces the bulk radius, thereby constraining the spectrum within a more stable regime (Fig 6C and D). Although this approach does not preserve the original dynamical structure as well, it may still retain partial low-dimensional dynamics while maintaining stability.

**Fig 6.**
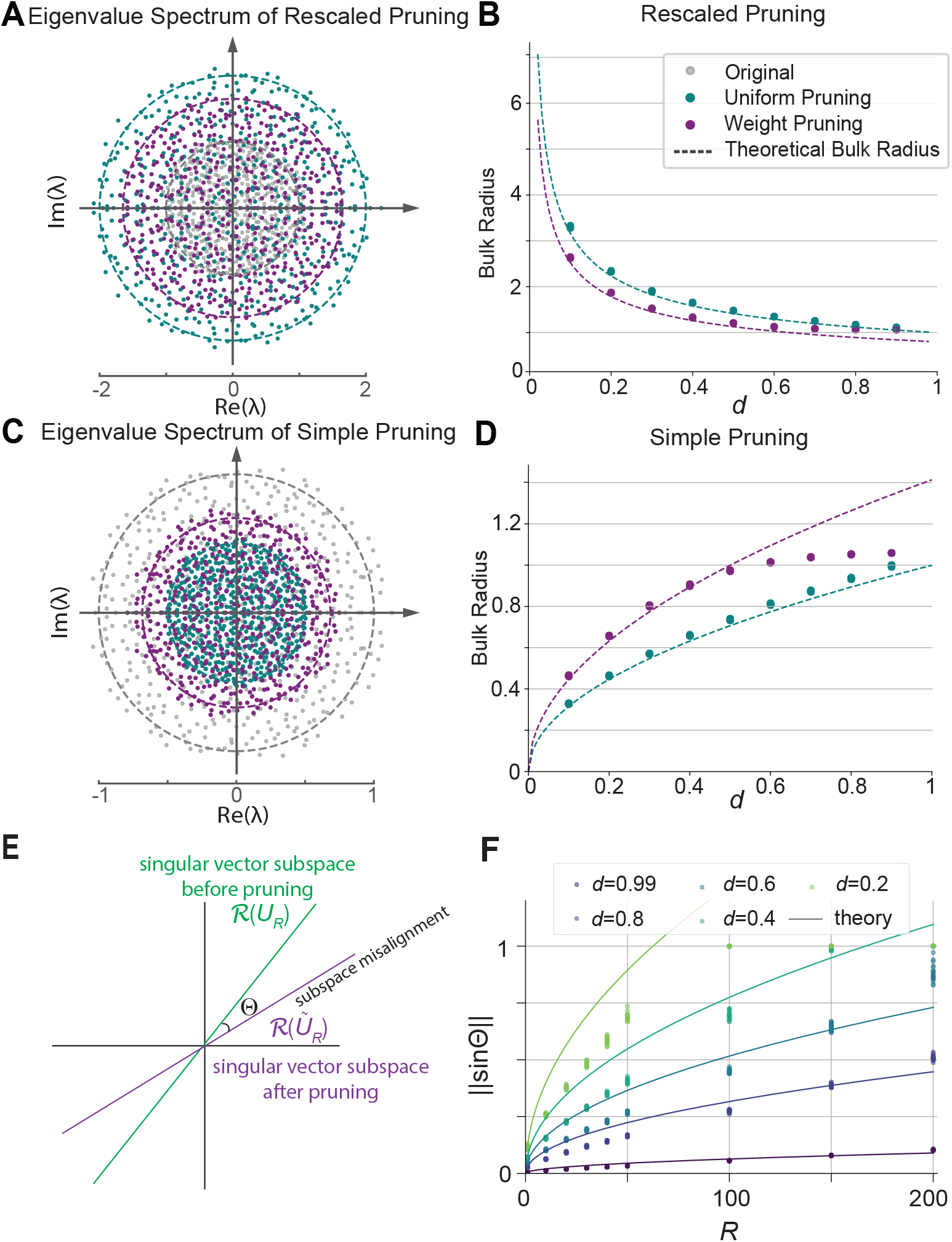
Eigenvalue spectrum of pruned full-rank RNNs and singular vector subspace rotations caused by pruning high-rank RNNs. **A**: Distribution of eigenvalues in the complex plane for pruned full-rank RNNs with rescaling. Turquoise dots indicate uniform pruning and purple dots weight pruning. The corresponding dashed lines indicate the theoretical bulk radii. **B**: Dependence of the bulk radius on density *d* under rescaled pruning. Turquoise and purple represent uniform and weight pruning, respectively. Dots represent the numerical bulk radii, computed by averaging the 50 largest eigenvalues (7 trials per condition), while dashed lines represent theoretical estimation. **C, D**: Same as panels **A** and **B**, but under simple pruning. For weight pruning, the theoretical estimation of the bulk radius is accurate only at low densities *d*, since the theoretical estimation assumes *p*_*ij*_ = *K*|*J*_*ij*_| ≤1, which holds only in the low-density regime. **E**: Schematic of subspace rotations under pruning. The subspace of singular vectors before pruning is represented by a green line and that after pruning is represented by a purple line. Θ represents the misalignment angle. **F**: Theoretical and numerically-observed rotations under uniform pruning for various ranks and pruning densities. Solid lines indicate theoretical upper bounds (Eq 12). Dots represent values from numerical simulations (20 trials per condition). *N* = 1000 across all panels.

Overall, for high rank RNNs, rescaled pruning gradually leads the network to instability as the connectivity density *d* decreases, while simple pruning preserves network stability across all densities.

#### Misalignment of Singular Vectors

Preserving the eigenvalue spectrum alone is not sufficient to maintain network functionality, as network dynamics also depends on the left and right singular vectors. In rank-one networks, network dynamics depend on the alignment of the input vector with the right connectivity vector ***n***, and on that of the readout vector and the left connectivity vector ***m*** [12]. For a higher-rank connectivity matrix as in Eq 3, we may replace ***m*** and ***n*** in the previous statement by the subspaces spanned by 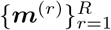 and 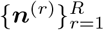, respectively. In this section, we examine the rotations of the subspaces spanned by the left and right singular vectors under uniform pruning.

The SVD of original connectivity matrix ***J*** ∈ ℝ^*N*×*N*^ of rank *R* is given by ***J*** = ***U* Σ*V*** ^⊤^, where **Σ** contains *R* non-zero singular values. The connectivity matrix after pruning, 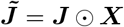, is formally full-rank and may be decomposed as

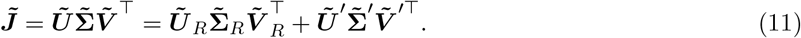

Here 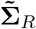, ***Ũ***_*R*_, and 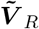 contain the leading *R* singular values, right and left singular vectors, and 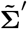, ***Ũ***^′^, and 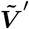 the trailing *N* − *R* ones, respectively.

We are interested in the misalignment between the original and pruned subspace spanned by top *R* singular vectors (Fig 6E). This is quantified via the canonical angles between the subspaces, which we denote Θ(ℛ (***U*** _*R*_), ℛ(***Ũ***_*R*_)) and 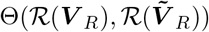. By symmetry, the left- and right-singular vectors are statistically identical, so we only consider Θ(ℛ (***U*** _*R*_), ℛ (***Ũ***_*R*_)) ≡ Θ.

Wedin’s theorem provides a way to approximate the singular vector subspace rotations after a random perturbation [67]. Assuming simple uniform pruning, we may cast the effects of pruning as a zero-mean perturbation ***T***, i.e., 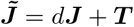 (where *d* is the expected density after pruning, defined in Eq 8). Then Wedin’s theorem gives an upper bound on the subspace rotation, ∥sin Θ∥ ≤ ∥***T*** ∥*/δ*, where *δ* = *dσ*^2^ (*σ* = *σ*_*m*_ = *σ*_*n*_) is the gap between the *R* and *R* + 1 singular values of the scaled matrix *d****J***, and ***T*** is the spectral norm of the pruning-induced perturbation. Using asymptotic arguments, we have for large *N* that 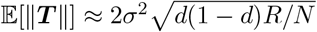, giving us the asymptotic bound

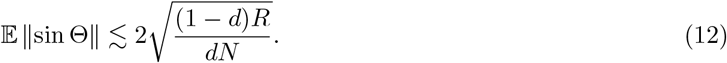

See Methods for a heuristic derivation.

Eq 12 agrees with values of ∥sin Θ∥ obtained from numerical simulations across various ranks *R* and density levels *d* (Fig 6F). These results indicate that low-rank matrices preserve their singular vector subspaces remarkably well after pruning. This gives further explanation of why network dynamics in low-rank networks are preserved with rescaling: the singular vectors are robust to pruning, so restoring the eigenvalue spectrum is sufficient to preserve the dynamics. On the other hand, with added ranks, the subspaces after pruning become increasingly misaligned with the original subspace. For high-rank networks, both the eigenvalue spectrum and the rotation of singular vectors must be considered to evaluate the preservation of network dynamics post pruning.

#### Identifying the Effects of Eigenspectra and Singular Vectors

We next examine how shifts in eigenvalues (Figs 1 and 6A-D) and singular vectors (Fig 6E and F) affect network dynamics. Specifically, we initialize a random rank-*R* connectivity matrix ***J*** according to Eq 3 and normalize it by the absolute value of the largest-magnitude eigenvalue as done in Fig 4. We define the input vector to be the top right singular vector of ***J*** . A random input current (duration 30 time units) is delivered via the input vector. The network firing rate ***r***(*t*) ≡ *ϕ*(***x***(*t*)) is recorded for 40 time units following stimulus onset. We then repeat the process after pruning ***J*** and calculate the MSE to measure the change in dynamics (see Methods for details). In addition, the rotation sin Θ of the left-singular vector subspace and the change in the magnitude of the largest eigenvalue induced by pruning of the connectivity matrix are measured. This setting allows us to directly measure the change in the rate dynamics and its relation to the eigenvalue spectrum and subspace rotations.

When uniform pruning with rescaling is applied (Fig 7A), we find that both eigenvalues and subspaces are preserved for low *R* but not for high *R*, as expected from previous results. The firing rate MSE increases with both *R* and sparsity, reflecting the combined effects of these factors and mirroring the results of the flip-flop task (Fig 5C and E, solid lines). With simple uniform pruning (Fig 7B), the leading eigenvalues decrease uniformly across ranks, reflecting the observations of Fig 1. On the other hand, subspace rotations are rank-dependent as in the rescaled uniform pruning, as rescaling does not affect subspace rotations. The eigenvalues have a dominant influence on the rate MSE, which is reflected in the strong *d*-dependence. This mirrors our results above, showing the distortion of the eigenvalue spectrum for low-rank matrices (Fig 1) and the inability of low-rank RNNs trained on the flip-flop task to preserve their dynamics after simple pruning (Fig 5 C and E, dashed lines).

**Fig 7.**
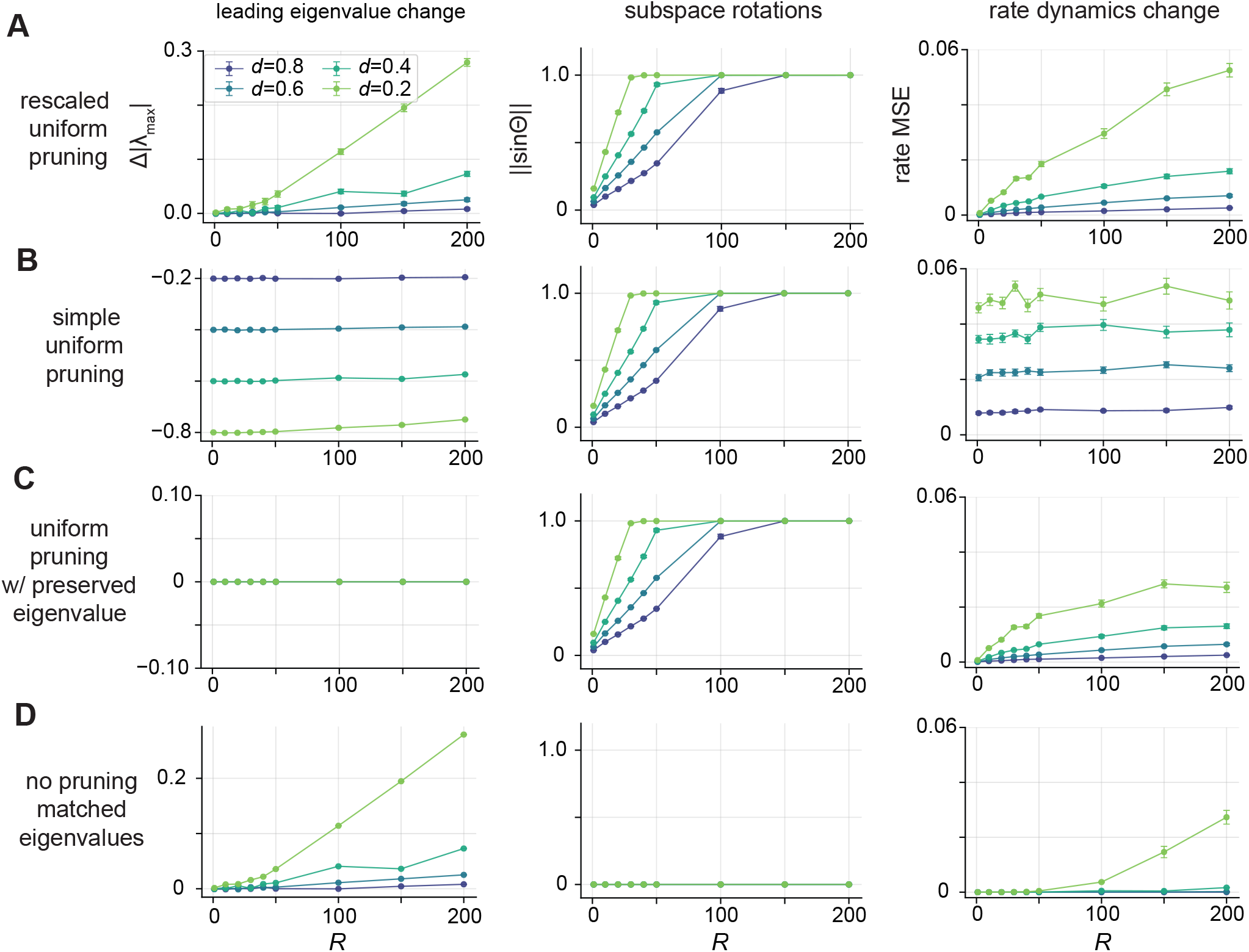
Isolating effects of eigenvalue and singular vector rotations on dynamics. **A**: Comparison of connectivity matrix properties and untrained RNN dynamics after rescaled uniform pruning. Shown are changes in the highest eigenvalue magnitude (left), subspace rotations (middle), and changes in network rate dynamics (right) in response to a pulse of random noise. **B**: Same as **A**, but for simple uniform pruning. **C**: Results for uniform pruning, but with the connectivity matrix scaled exactly to preserve the leading eigenvalue. **D**: Results in which the network was not pruned, but scaled to match the mean eigenvalue increase as seen in **A** from rescaled pruning. Across all panels, *N* = 500 and error bars indicate the mean ± sem from 20 simulations per condition.

To isolate the effects of subspace rotations, we perform uniform pruning, but rescale the connectivity to precisely maintain the leading eigenvalue by directly scaling by |*λ*_max_| (Fig 7C). This results in completely preserved leading eigenvalues, but pruning still induces subspace rotations. Now, the rate MSE reflects the rank-dependence of the subspace rotations, confirming the effect of singular vector rotations. Compared to the rescaled pruning case, the rate MSE is generally smaller, especially for high ranks when *d* = 0.2.

We then isolate the effects of eigenvalue inflation by scaling the connectivity matrix by a scalar value to match the inflation observed in Fig 7A, which inflates the eigenvalues only without inducing subspace rotations (Fig 7D). Again, we observe a rate MSE that reflects the eigenvalue inflation, but much smaller in magnitude than in Fig 7A. This confirms that the change in dynamics induced by uniform pruning comes from the effects of both eigenvalue inflation and singular vector subspace rotations.

### Rescaling is Critical in Low-Rank Pruning

Our analysis using the uniform and weight pruning rules highlights that rescaling is the key factor for preserving the eigenvalue spectrum in the low-rank regime, thereby maintaining the underlying dynamics. To draw a closer parallel to biological systems, we also explore pruning rules inspired by Hebbian learning, which posits that neurons that fire together tend to strengthen their synaptic connections [68–71].

Specifically, we examine three pruning rules,

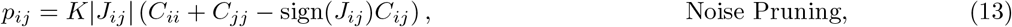

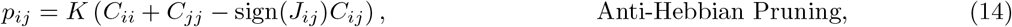

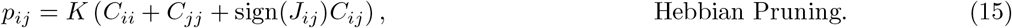

Noise pruning (Eq 13) is a biologically motivated pruning scheme in which the pruning probability depends on the covariance matrix ***C*** of neural activity under noisy input along with connection weights [45]. To isolate the contribution of neural activity correlations, we explore two variations of noise pruning (Eqs 14 and 15), in which the weight-dependent factor is removed, resulting in purely Hebbian and anti-Hebbian pruning rules [71]. We incorporate rescaling in all the pruning rules.

Each of these pruning rules relies on the covariance structure of network activity. Anti-Hebbian pruning preferentially removes connections between highly correlated neurons (large *C*_*ij*_), effectively decorrelating network activity. Hebbian pruning instead preserves such correlated connections while removing those between weakly correlated neurons. Noise pruning combines anti-Hebbian with weight pruning, probing whether observed correlations exceed expectations based on synaptic weight.

Despite their greater biological plausibility, these Hebbian-inspired pruning strategies produce qualitatively similar outcomes to our simpler pruning rules. Most notably, all these methods, when combined with rescaling, preserve the leading eigenvalue in low-rank matrices, thus maintain low-dimensional dynamics (Fig 8). This suggests that, functionally, these biologically motivated pruning rules induce the similar effect on dynamics of low-rank networks as uniform and weight pruning.

**Fig 8.**
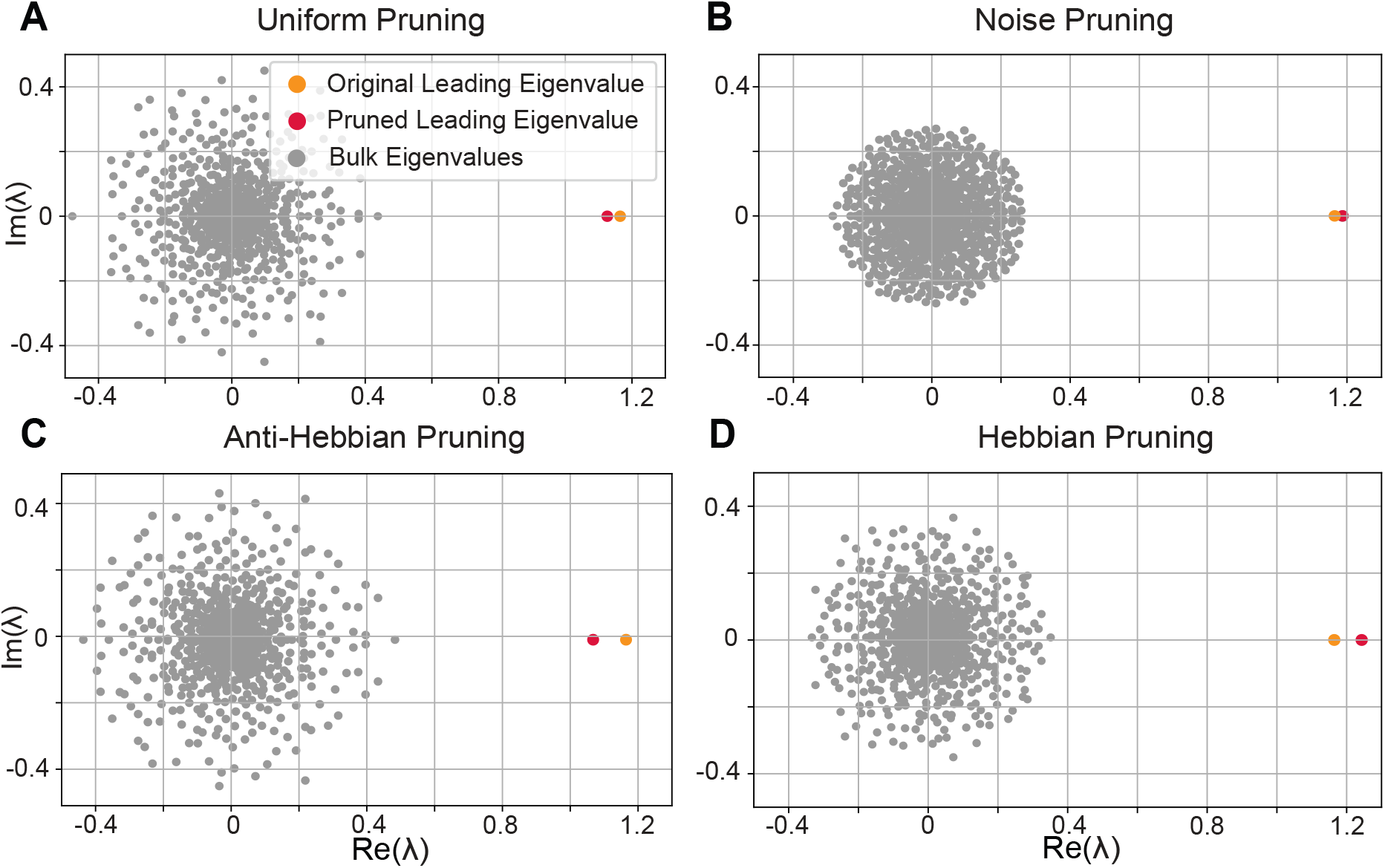
Effects of biologically inspired pruning rules on network eigenvalue spectrum. **A-D**: Eigenvalue spectrum of the connectivity after (**A**) uniform pruning, (**B**) noise pruning, (**C**) anti-Hebbian pruning, and (**D**) Hebbian pruning on a rank-one RNN. Each rule prunes the RNN to the density *d* = 0.1. For all panels, *N* = 1000 and rescaling was applied. All methods, when combined with rescaling, preserve the leading eigenvalue.

More importantly, it suggests that in the low-rank regime, rescaling alone is often sufficient to preserve the eigenvalue spectrum and maintain the low-dimensional dynamics. This finding highlights a key insight: the low-rank structure of the network is inherently robust to pruning whenever rescaling is applied, emphasizing that synaptic scaling to maintain network homeostasis is a powerful and potentially sufficient mechanism for preserving the network’s core dynamics under pruning. This cannot be generalized the to high-rank regime, as their dynamics are not dominated by a single leading eigenvalue and subspace rotations are substantial, making rescaling not appropriate to preserve the low-dimensional dynamics.

## Discussion

In this study, we examine how different pruning schemes affect the low-dimensional dynamics of nonlinear RNNs, with particular emphasis on the role of network rank. We find that the low-dimensional dynamics and the computational capabilities of an RNN can be preserved if the network is of low-rank and pruning is rescaled. In this regime, rescaling maintains the leading eigenvalue of the connectivity matrix after pruning, thereby preserving the geometry of low-dimensional trajectories. In contrast, simple pruning without rescaling consistently diminishes the leading eigenvalue, distorting the dynamics even when the random bulk remains small [20]. In high-rank networks, however, rescaling does not prevent the degradation of dynamics followed by pruning, as increasing the rank leads to greater expansion of the bulk spectrum and pruning-induced misalignment of subspaces. These results reveal a critical rank dependency for the performance of rescaled pruning, suggesting that low-rank connectivity exhibits inherent robustness against pruning and supports stable computation under biologically realistic sparsity constraints.

From a biological perspective, our rescaling approach captures homeostatic synaptic scaling, a regulatory process in which neurons globally adjust synaptic strengths to stabilize firing rates following changes in connectivity [46–60]. By ensuring that the population-averaged connectivity weight remains constant post pruning, the synaptic rescaling effectively stabilizes the network’s average firing rates. Such stabilization is essential for probabilistic pruning in the low-rank regime, regardless of the specific pruning method employed. While our primary pruning rules are chosen to keep our mathematical analysis simple, Hebbian-inspired variants produce qualitatively similar results where rescaling preserves the leading eigenvalue locations post pruning in low-rank networks. This indicates that the preservation of dynamics depends more on synaptic rescaling than on the exact pruning rule for selecting synapses to eliminate.

Conceptually, rescaled pruning shares similarity with Hebbian plasticity under a fixed sparsity constraint, in that both modify synaptic weights while maintaining a constrained connectivity structure. Prior works have employed Hebbian rules to continuously update synaptic weights based on activity correlations under a fixed sparse mask [72, 73]. However, rescaled pruning is distinct from such activity-dependent pruning rules in two important ways. First, activity-dependent pruning operates on a fixed sparse connectivity mask and subsequently updates synaptic strengths through activity-dependent plasticity, whereas in rescaled pruning the sparsification and synaptic modifications occur simultaneously when the pruning mask is applied. Second, rescaled pruning depends only on the synaptic weights themselves rather than on pre- and postsynaptic neuronal activity. As a result, synapses are either eliminated or rescaled to preserve the average connectivity strength across the network.

While some works [50] model homeostatic synaptic scaling to depend explicitly on neural activity, other studies have also implemented homeostatic scaling by fixing the total weights [61], similar to our implementation of synaptic rescaling. Indeed, this study [61] showed that their generative model with a fixed net weight replicates realistic connectivity structures in fly mushroom body, suggesting the biological plausibility of this implementation of homeostatic synaptic scaling.

Our findings reveal a pruning strategy that preserves low-dimensional dynamics in low-rank RNNs, offering potential insight into the organization of biological networks. Although low-rank network theory has produced elegant analytical advances, there is no direct evidence that cortical networks are inherently low-rank, despite recent efforts to infer global low-rank connectivity from low-dimensional local neuron population statistics [74, 75]. Our findings suggest an alternative hypothesis: biological neural circuits such as cortical networks may initially inherit low-rank connectivity and subsequently undergo developmental pruning, resulting in sparse, formally full-rank networks that remain effectively low-rank regarding their dynamics. Such pruning does not produce a sharp cutoff in the singular value spectrum. Instead, the spectrum decays smoothly with a rapid falloff, a property consistent with observations in the Drosophila hemibrain [76] as well as in other complex systems such as *C. elegans* and human gut microbiome networks [77–79].

It is also worth noting that, beyond the obvious benefits of reducing metabolic and wiring costs [34–36], the functional advantages of sparsity remain unclear in biological systems. While our work shows that rescaled pruning can maintain network dynamics and computational performance under certain conditions, whether the observed sparsity in biological neuronal networks enhances the brain’s functional capacity, remains unclear. However, results from deep learning suggest that networks could sometimes benefit from high levels of sparsity with marginal loss of accuracy [80, 81], which can act as a natural regularizer to mitigate over-parameterization and overfitting [82]. This raises the possibility that biological sparsity may serve a similar dual role, conserving resources while supporting stable, generalizable computation.

Altogether, our results highlight the interplay between network rank, sparsity, and synaptic rescaling in preserving the low-dimensional dynamics in RNNs. By bridging theoretical low-rank RNN models with biologically plausible pruning mechanisms, we provide a framework for understanding how neural circuits may retain computational capacity despite extensive synaptic loss during development and across the lifespan resulting in their sparse connectivity.

## Methods

### Simulation Details

#### Flip-Flop Task

Each trial of duration *T* begins with a fixation period of length *t*_fix_ with no input, followed by a sequence of brief stimulus pulses. Each pulse lasts for *t*_stim_ and activates a single input channel *k* with value *u*_*k*_(*t*) = *su*_amp_, while all other inputs remain zero. The channel *k* and sign *s* ∈{± 1} are chosen randomly for each pulse.

Each stimulus is followed by a decision period of duration *t*_decision_, after which the choice period begins and the loss function is activated. During the choice period, the target output is set to 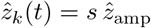 with all other outputs at zero. The loss is inactive outside of the choice periods. A choice period ends with the onset of the next stimulus pulse. The inter-stimulus interval *T*_interval_ is drawn randomly.

Here we set 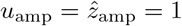, *t*_fix_ = 5, *t*_stim_ = 1, *t*_decision_ = 5, *T*_interval_ ∼ 𝒰 (17.5, 25), and *T* = 150.

#### RNN Training

In this paper, we consider two training schematics for RNNs: pre-pruning training and post-pruning retraining. Both approaches aim to train the RNN to perform the flip-flop task using the gradient-based optimizer *Adam*.

In pre-pruning training, we apply rank-constrained backpropagation (Algorithm 1) to the input vectors ***I***^(*k*)^, recurrent connectivity ***J***, and output weights ***w***^(*l*)^. In contrast, post-pruning retraining updates only the output weights ***w***^(*l*)^, with the goal of restoring task performance after pruning a pre-trained RNN.

For gradient-based optimization, we define the quadratic loss function

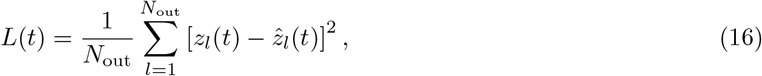

where *z*_*l*_(*t*) denotes the model readout, 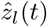 denotes the target output, and *N*_out_ denotes the number of output channels.

The learning rate is set to *η* = 0.0001 for single- and double-channel flip-flop tasks, and *η* = 0.01 for post-pruning retraining. Networks are trained for 200 epochs on the flip-flop task, where each epoch consists of 10 batches of 100 independent trials. Each trial contains 6–7 input signals. For readout retraining, the network is trained for 100 epochs with the same number of trials.

All the RNN training is done with PyTorch (v2.10.0).

#### General Setup for RNN Dynamics and Spectral Analysis

The network dynamics, as described in Eq 1, is simulated using the forward Euler scheme, with time constant *τ* = 1 and Δ*t* = 0.1 for random RNNs (Fig 2, Fig 4 and Fig 7) and Δ*t* = 0.5 for task-trained RNNs (Fig 3 and Fig 5).

For empirical analyses of the leading eigenvalue and bulk radius, we use networks of size *N* = 1000 (Fig 1, Fig 6A–D and Fig 8). For untrained RNNs, we set *N* = 800 (Fig 2, Fig 4 and Fig 7). For task-trained RNNs, *N* = 500 (Fig 3 and Fig 5B–E). Results are robust to the choice of *N* as long as it is sufficiently large.

For task-trained RNNs, the initial condition is ***x***(0) = **0**. For random RNNs (Fig 2 and Fig 4B–E), initial conditions are drawn independently from a uniform distribution *x*_*i*_(0) ∼ 𝒰 (−0.01, 0.01). For Fig 7, the initial condition is ***x***(0) = **0**.

For simulations of high-rank random (untrained) RNNs (Fig 4, Fig 6E–F and Fig 7), we construct ***J*** for a given *R* directly from Eq 3 with *σ*_*m*_ = *σ*_*n*_ = *σ*_*mn*_ = 1. Because we examine *R* up to 500, we no longer have the condition *R* ≪ *N* satisfied, and the scaling of Eq 3 does not preserve the spectral radius independent of *R*. We therefore normalize each generated *J* by the empirical leading eigenvalue to ensure that the leading eigenvalue remains independent of *R*.

Simulations in Fig 2, Fig 4 and Fig 7 are implemented in Python (v3.12.13).

#### Input-Driven RNN Dynamics

For the simulations shown in Figs 2, 4 and 7, the RNN dynamics follows the equation

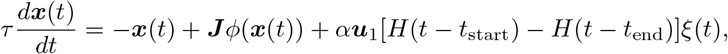

where *H* is the Heaviside step function, ***u***_1_ is the top right singular vector, *α* is a scalar parameter, *ξ*(*t*) is an input modulation function whose form depends on the simulation setting, and *t*_start_ and *t*_end_ determine the start and end of the stimulus, respectively.

In Fig 2, *ξ*(*t*) ≡ 1 and *α* = 1. The stimulus lasts for 60 time units with the stimulus onset and offset at *t*_start_ = 20 and *t*_end_ = 80, respectively. In Fig 4, *ξ*(*t*) ≡ 1 and *α* = 10. The stimulus duration is 60 time units with *t*_start_ = 30 and *t*_end_ = 90.

In Fig 7, *ξ*(*t*) is standard independent white noise with mean 0 and variance 1, and *α* = 50. The stimulus lasts for 30 time units with *t*_start_ = 10 and *t*_end_ = 40. The mean square error of the firing rates during times 10 ≤ *t <* 50 (400 time steps) before and after pruning (or scaling in the case of Fig 7D) is computed as

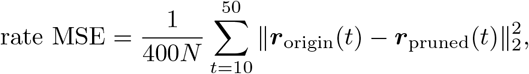

where ***r***_origin_(*t*) ≡*ϕ*(***x***_origin_(*t*)), ***r***_pruned_(*t*) ≡ *ϕ*(***x***_pruned_(*t*)) and the summation is in increments of the time step Δ*t*. This is repeated 20 times for each condition.

#### Pruning Parameter Selection

The post-pruning connection density *d* (Eq 8) depends on the control parameters 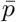 and *K* for uniform and weight pruning, respectively. To target a specific *d*, in the case of uniform pruning we simply set 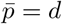. For weight pruning, because the mapping between *K* and density *d* depends on the specific connectivity matrix ***J***, the value of *K* required to achieve a target *d* is determined empirically. Since *d* increases monotonically with *K*, we use a binary search over *K*, initialized with the lower bound *K*_low_ = 1 and an upper bound *K*_high_ = 1000, and iteratively refine *K* until the resulting density *d* matches the target within a tolerance of 10^−4^.

### Exact scaling of rank-*R* matrices

Eq 3, as done in [12], describes low-rank matrices in the *R* ≪ *N* limit. If ***m*** and ***n*** are generated according to the bivariate Gaussian distribution (Eq 9), then the leading eigenvalue stays approximately fixed at *σ*_*mn*_ with respect to *R*. However, when the condition *R* ≪ *N* does not hold, random overlaps between the independent vectors ***m***^(*r*)^ will cause the leading eigenvalue to exceed *σ*_*mn*_. For this reason, in Figs 4 and 7, where we examine ranks up to *R/N* = 5*/*8 and 1*/*2, respectively, we normalize the matrix numerically by dividing by the largest empirical eigenvalue of each generated (pre-pruning) matrix. This ensures that the initial RNN stays within a stable dynamical regime across *R*. Below we provide results on eigenvalue scaling with *R* for two special cases, *σ*_*mn*_ = 0 (no correlation) and *σ*_*mn*_ = *σ*_*m*_ = *σ*_*n*_ (completely correlated).

#### Maximum-eigenvalue-preserving scaling rule for *σ*_*mn*_ = 0

When *σ*_*mn*_ = 0, we may write

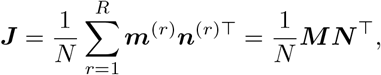

where ***M*** and ***N*** are *N* × *R* matrices whose *r*th column is ***m***^(*r*)^ and ***n***^(*r*)^, respectively. ***J*** thus has the same non-zero eigenvalues as the *R* × *R* matrix ***Q*** = *N* ^−1^***N*** ^⊤^***M*** . This matrix has elements

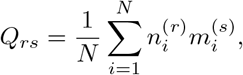

which have mean zero and variance

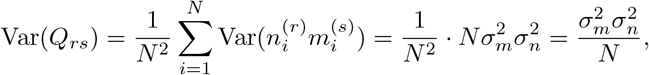

due to the independence of 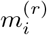 and 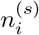. We then invoke Girko’s circular law, which says that the eigenspectrum of ***Q*** converges to a uniform distribution over a disk with radius 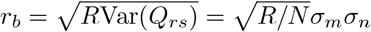 for large *R*. Thus to remove the growth of the leading eigenvalue with increasing *R*, one may choose the scaling

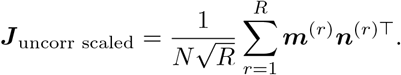

#### Maximum-eigenvalue-preserving scaling rule for *σ*_*mn*_ = *σ*_*m*_ = *σ*_*n*_

In this case we have ***m***^(*r*)^ = ***n***^(*r*)^ and

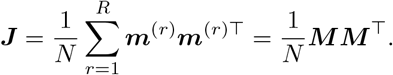

This is a Wishart matrix [83], whose eigenvalues follow the Marchenko-Pastur distribution [84] whose support is 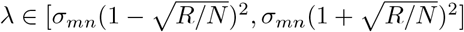. Hence the leading eigenvalue scales as 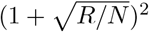. The scaling rule that preserves the leading eigenvalue in this case would be

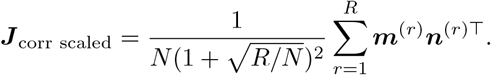

### Derivation of Eigenvalue Spectrum of Rank-one Matrix

A rank-one connectivity matrix is defined as ***J*** = *N* ^−1^***mn***^⊤^, where the vectors ***m, n*** ∈ ℝ^*N*^ are drawn from a joint Gaussian distribution characterized by

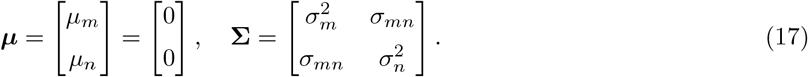

The sparse matrix 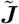 is generated by applying a mask to ***J***, namely 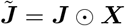. In simple pruning

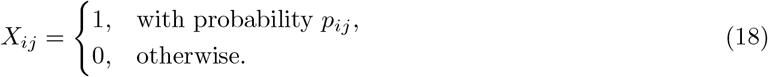

In rescaled pruning,

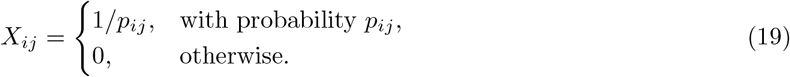

Note that *p*_*ij*_ represents the probability of preserving a connectivity edge. Namely,

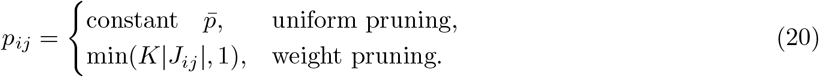

#### Leading eigenvalue

We look at the product of 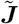 and ***m***, where ***m*** is the original eigenvector of ***J*** that corresponds to the original leading eigenvalue 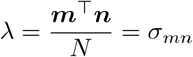 in the large *N* limit.

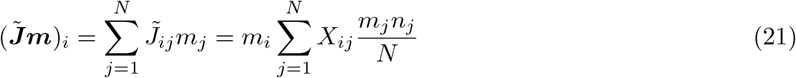

As *N* → ∞, due to the law of large numbers, we have

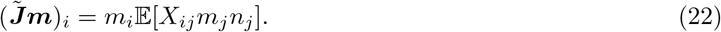

This makes ***m*** an eigenvector and 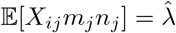 the associated eigenvalue, which is the leading eigenvalue of the sparse matrix 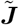. We proceed to compute the eigenvalue for each pruning rule.

In the uniform pruning case, *X*_*ij*_ is independent of *m*_*j*_ and *n*_*j*_, so we have

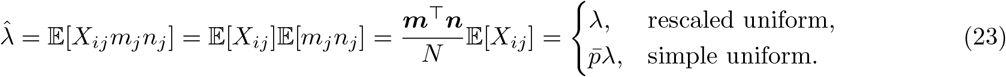

For weight pruning, we need to estimate 𝔼[*X*_*ij*_*m*_*j*_*n*_*j*_] without assuming independence.

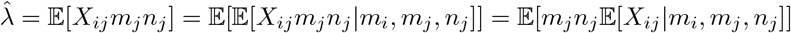

For rescaled weight pruning, we have 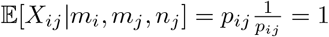, thus

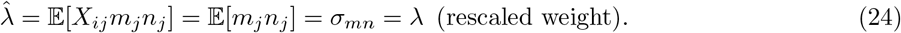

For simple weight pruning, we have 𝔼[*X*_*ij*_|*m*_*i*_, *m*_*j*_, *n*_*j*_] = *p*_*ij*_ = min(*K*|*J*_*ij*_|, 1). We use the approximation *p*_*ij*_ = *K*|*J*_*ij*_|. This will lead to an overestimation of *p*_*ij*_ when *K* is high, as high values of *K* will increase the probability that *K*|*J*_*ij*_| is larger than 1. Using this estimation, we have

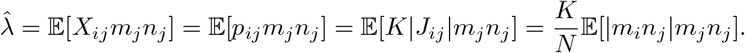

We recall that *m*_*i*_ and *n*_*j*_ are independent when *i* ≠ *j*, and correlated when *i* = *j*. Here, we neglect the diagonal elements in *J*, where *i* = *j*, as they contribute only an *O*(1*/N* ) fraction and are negligible in the large *N* limit. Thus,

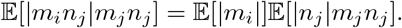

Taking the marginal distribution, we have 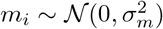 and so 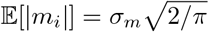. We also have

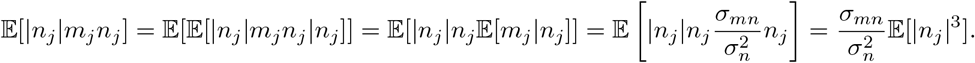

Here we use the standard identity that given a normally-distributed random variable *Z* ∼ 𝒩 (0, *σ*^2^), for *k >* 0,

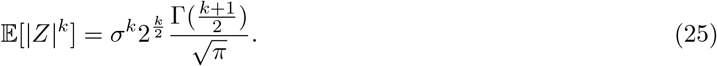

Using *k* = 3, Γ(2) = 1, we have 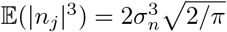. Thus

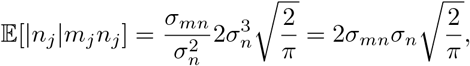

and we finally have

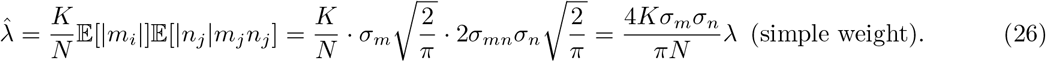

In summary, the leading eigenvalues after weight pruning are

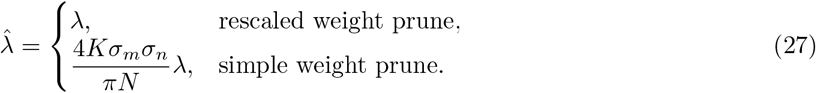

We have shown that in the large *N* limit, ***m*** remains an eigenvector under different pruning rules. Additionally, if 𝔼[*X*_*ij*_ | *m*_*i*_, *m*_*j*_, *n*_*j*_] = 1, which represents rescaling, the corresponding eigenvalue will always be preserved.

#### Bulk radius

We first introduce the circular law. Assume that a matrix ***A*** ∈ ℝ^*N*×*N*^ is drawn from distribution with 𝔼[*A*_*ij*_] = 0. Then as *N* → ∞, the eigenvalue spectral distribution converges to the uniform distribution over a disk of radius 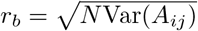 [62, 63].

To determine the radius of the bulk distribution of eigenvalues, we derive the variance of the elements of the matrix with the outlier eigenvalue removed 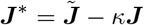,where *κ* is the scaling factor of leading eigenvalue 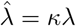. For simplicity, here we let *σ* = *σ*_*m*_ = *σ*_*n*_. Using the fact that 𝔼[*X*_*ij*_] = 1 and 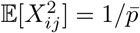 for rescaled uniform pruning and 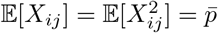 for simple uniform pruning, we have

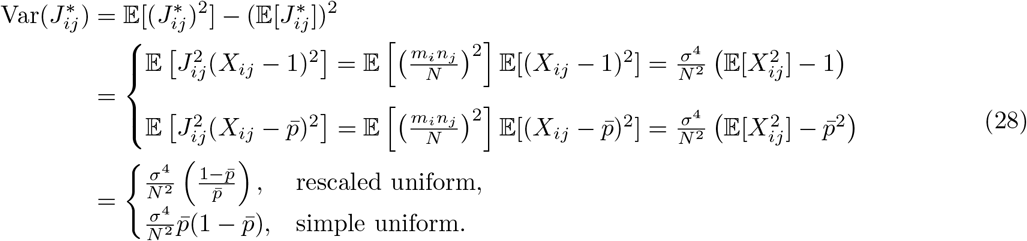

This gives a bulk radius of 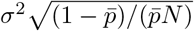 for rescaled uniform pruning and 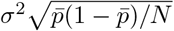 for simple uniform pruning.

For weight pruning, the estimate follows a similar strategy. Let *η* = 4*Kσ*^2^*/πN* . From the previous section, for simple weight pruning, the leading eigenvalue is given as 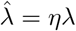. Then,

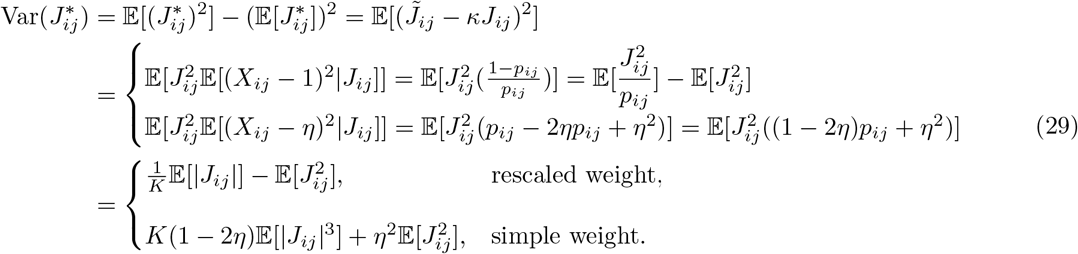

When estimating the moments of |*J*_*ij*_|, we still assume *i* ≠ *j*, as diagonal elements are negligible for large *N* . For the non-diagonal case, *m*_*i*_ and *n*_*j*_ are independent of each other and *m*_*i*_ ∼ 𝒩 (0, *σ*_*m*_), *n*_*j*_ ∼ 𝒩(0, *σ*_*n*_). Using Eq 25, we have

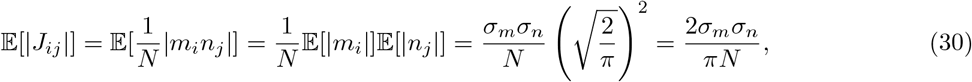

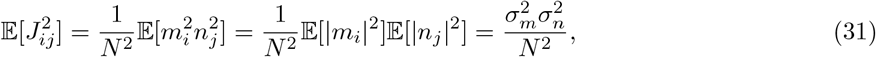

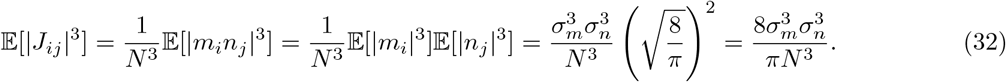

Substituting the above expectations into Eq 29 and using 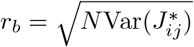, we have

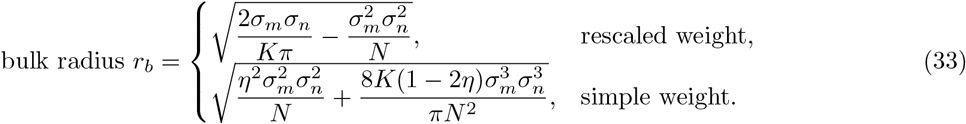

While we neglecte diagonal elements in the derivations above, it is also worth noting that one can obtain a better estimation of moments of |*J*_*ij*_| by averaging over diagonal and non-diagonal elements. The result incorporating diagonal elements are given as follows, although we omit them in our calculation for simplicity: with the given distribution of ***J***, we have 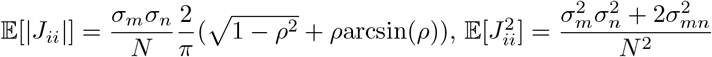 and 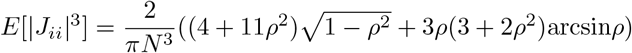 where 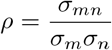 [85] [86].

### Derivation of Eigenvalue spectrum of Full-rank Matrix

#### Uniform pruning

We assume *J*_*ij*_ ∼ 𝒩(0, *σ*^2^*/N* ) and the pruned connectivity matrix is given by 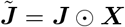, where the elements of the masking matrix ***X*** follow a Bernoulli distribution, specifically 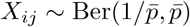 for rescaled pruning and 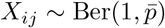 for simple pruning.

Since *J*_*ij*_ is symmetrically distributed around zero and *X*_*ij*_ only depends on |*J*_*ij*_|, the product *J*_*ij*_*X*_*ij*_ remains symmetrically distributed around zero, thus 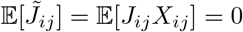. This holds for both rescaled pruning and simple pruning. By this we have,

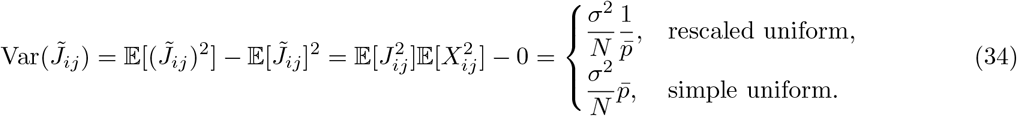

where we use the independence of *J*_*ij*_ and *X*_*ij*_ in the second step. Thus, rescaling will lead to a bulk radius of 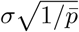, while the bulk radius is 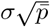 without rescaling.

#### Weight pruning

For weight pruning we still have 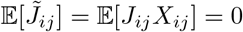. However, the masking matrix depends on ***J*** here, which requires a different calculation. Again, with the assumption of sufficiently small *K*, we make the approximation *p*_*ij*_ = *K*|*J*_*ij*_|, and have

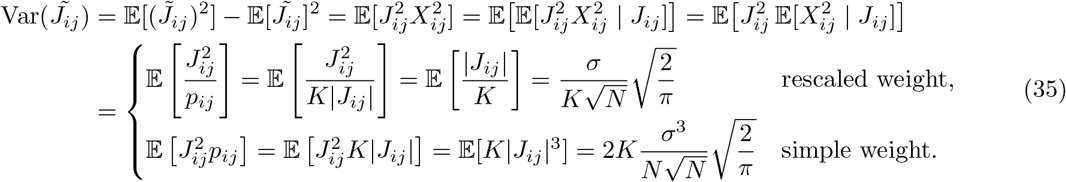

This gives a bulk radius of 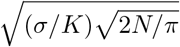 for rescaled pruning and 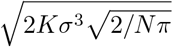 for simple pruning.

### Subspace rotations under uniform pruning

Here we provide a heuristic derivation of subspace rotation bound of Eq 12 under uniform pruning. We assume that an initial connectivity matrix of rank *R* is constructed according to Eqs 3 and 9, where we set *σ*_*m*_ = *σ*_*n*_ = *σ*. Note that because the entries of ***m***^(*r*)^ and ***n***^(*r*)^ have variance *σ*^2^, their norms have an expected value of 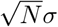. Ignoring overlaps between modes *r* (which vanish for large *N* ), each nodes contribute a singular value of *N* ^−1^ ∥***m***^(*r*)^ ∥ ∥***n***^(*r*)^ ∥ ≈ *σ*^2^. Since there are *R* such rank-one matrices, ***J*** will have *R* singular values at *σ*^2^ in expectation and the remaining *N* − *R* singular values will be 0. Hence there will be a gap of *σ*^2^ between the *R*-th and *R* + 1-th singular values.

We first consider simple uniform pruning. The connectivity matrix after pruning can be written

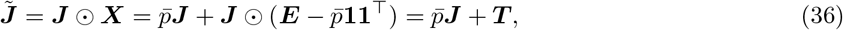

where 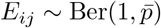 and **11**^⊤^ is a square matrix whose elements are 1. This gives 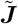 as a sum of a scaled version of the original matrix that represents the expectation under pruning, i.e. 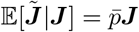, and a zero-mean perturbation ***T*** such that 𝔼[*T*_*ij*_] = 0.

We now state Wedin’s Theorem [67] in its original form, then a more specialized corollary applicable to our case.

**Theorem** (Wedin’s sin *θ*-theorem) Let ***A*** be a complex *m* × *n*-matrix with singular value decomposition

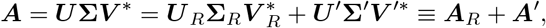

where ***V*** _*R*_ (or ***U*** _*R*_) contains the first *R* left (right) singular vectors and ***V*** ^′^ (***U*** ^′^) the next *R*^′^, such that the rank of ***A*** is *R* + *R*^′^. **Σ**_*R*_ is a diagonal matrix with diagonal entries *σ*_1_, *σ*_2_, · · ·, *σ*_*R*_ and **Σ**^′^ is one with entries *σ*_*R*+1_, · · ·, *σ*_*R*+*R*_*′*, where the singular values are ordered such that *σ*_1_ ≥ *σ*_2_ ≥ · · · ≥ *σ*_*R*_. Let ***B*** = ***A*** + ***T*** be the perturbation of ***A*** with a corresponding SVD. Let ***Y*** _*R*_ = [***y***_1_, · · ·, ***y***_*R*_], where ***y***_1_, · · ·, ***y***_*R*_ are orthonormal vectors spanning the subspace ℛ (***B***_*R*_) and analogously define ***X***_*R*_ through orthonormal vectors spanning 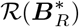. Define 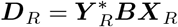. Define the residuals

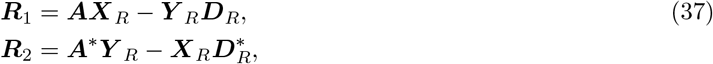

and take *ϵ* = max(||***R***_1_||, ||***R***_2_||).

Assume there exists an *α* ≥ 0 and *δ >* 0 such that *σ*_min_(***B***_*R*_) ≥ *α* + *δ* and *σ*_max_(***A*′**) ≤ *α*. Then

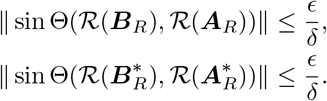

#### Corollary

It is the case that

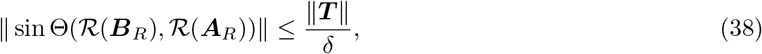

and similarly for sin 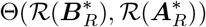.

*Proof*. Rewrite the residuals as 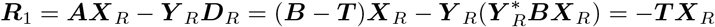 and similarly ***R***_2_ = −***T*** ^∗^***Y*** _*R*_. Then we have ∥***R***_1_∥ ≤ ∥***T***∥∥***X***_*R*_∥ ≤ ∥***T***∥ and similarly ∥***R***_2_∥ ≤ ∥***T***∥ . Eq 38 then immediately follows from Eq 37.

We thus apply Wedin’s Theorem to 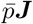 and 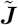. Note that since, as noted above, the construction under Eq 3 yields a gap of *σ*^2^ between the *R* and *R* + 1-highest singular values, 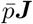 has a gap of 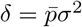. So Wedin’s Theorem yields

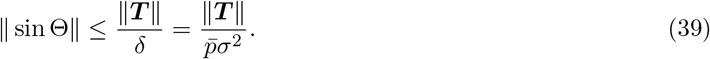

We evaluate the asymptotic form of ∥***T*** ∥ by first noting that for *i* ≠ *j*,

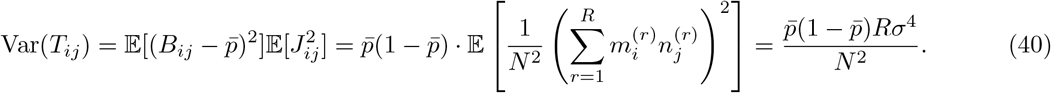

We base our argument on the variance of the off-diagonal elements only for simplicity. For completeness, we note that diagonal elements will have an additional dependence on *σ*_*mn*_. Noting that 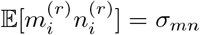 and 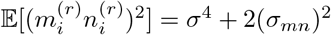, we have

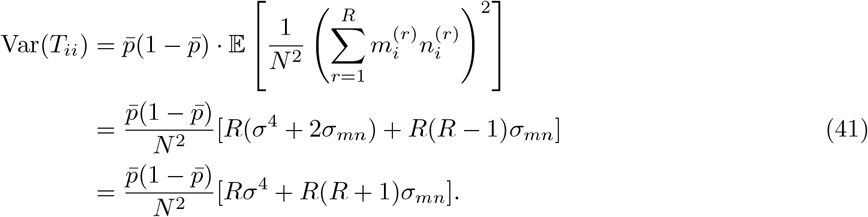

The Bai-Yin law [62, 87] states that a matrix ***G*** with i.i.d entries 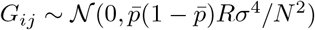 satisfies

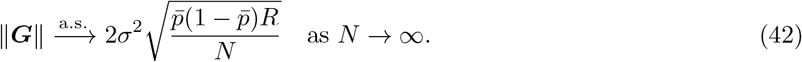

We conjecture that the norm of ***T***, whose off-diagonal entries match those of ***G*** up to the second moment, has the same asymptotic form. Plugging this into Eq 39 gives us the desired approximate bound of Eq 12. For the rescaled uniform pruning case, Eq 36 becomes 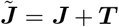 with 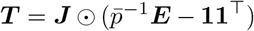. The result is that the perturbation norm becomes 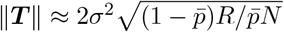 and *δ* becomes *σ*^2^, resulting in the same bound. Using the fact that 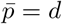 for uniform pruning, we finally have

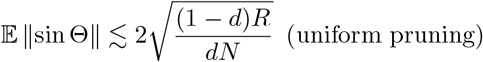

for large *N* .

## Code availability

Source code for all figures will be available upon publication on this Github repository: https://github.com/HChoiLab/Rescaled_Prunisng

## Notes

### Competing Interest Statement

The authors have declared no competing interest.

### Funding

- Alfred P. Sloan Foundation

